# A single cell RNA sequence atlas of the early *Drosophila* larval eye

**DOI:** 10.1101/2024.03.06.583750

**Authors:** Komal Kumar Bollepogu Raja, Kelvin Yeung, Yumei Li, Rui Chen, Graeme Mardon

## Abstract

The *Drosophila* eye has been an important model to understand principles of differentiation, proliferation, apoptosis and tissue morphogenesis. However, a single cell RNA sequence resource that captures gene expression dynamics from the initiation of differentiation to the specification of different cell types in the larval eye disc is lacking. Here, we report transcriptomic data from 13,000 cells that cover six developmental stages of the larval eye. Our data show cell clusters that correspond to all major cell types present in the eye disc ranging from the initiation of the morphogenetic furrow to the differentiation of each photoreceptor cell type as well as early cone cells. We identify dozens of cell type-specific genes whose function in different aspects of eye development have not been reported. These single cell data will greatly aid research groups studying different aspects of early eye development and will facilitate a deeper understanding of the larval eye as a model system.

## Introduction

Single cell technologies provide a powerful approach to characterize transcriptional and epigenomic states of single cells in multicellular organisms and have been transforming our understanding of biological tissues both in development as well as in disease (Kolodziejczyk, Kim et al. 2015, Wen, Li et al. 2022). Single cell technologies have been used in both basic and clinical research to investigate cellular heterogeneity, identify novel cell types, as well as to understand tumor ecosystems (Shapiro, Biezuner et al. 2013, Baslan and Hicks 2017, Lim, Lin et al. 2020). Therefore, generating comprehensive single cell omics atlases from model organisms will improve our understanding of cellular heterogeneity, cell fate determination, and tissue morphogenesis. Single cell atlases have been generated from tissues of several model organisms, including *mus musculus* (Consortium 2018), *Caenorhabditis elegans* (Taylor, Santpere et al. 2019) and efforts to generate comprehensive whole-organism single cell atlases across development are underway to understand organismal development and biology.

The model organism *Drosophila melanogaster* has been extensively studied for decades to understand conserved biological mechanisms underlying development, aging, neurodegeneration, among many other processes (Prüßing, Voigt et al. 2013, Yamaguchi and Yoshida 2018). Furthermore, the availability of sophisticated genetic tools, short generation times and a high degree of conservation with human genes make it a valuable resource for new discoveries. In addition, single cell omics methods have been applied to several *Drosophila* tissues (Karaiskos, Wahle et al. 2017, Davie, Janssens et al. 2018, Özel, Simon et al. 2021, Yeung, Bollepogu Raja et al. 2022, Bollepogu Raja, Yeung et al. 2023) and have led to numerous findings, including the discovery of new cell types. The *Drosophila* eye has been extensively used to decipher mechanisms of cellular morphogenesis, survival and differentiation. The adult *Drosophila* eye is a hexagonal lattice consisting of ∼750 repeating facets called ommatidia. Each ommatidium consists of eight neuronal and twelve non-neuronal cells. The eye emerges from a neuro-epithelial sac, the eye imaginal disc, and grows in size during larval stages. Cell type differentiation begins at the posterior equatorial margin of the late second instar eye disc with the initiation of a wave-like indentation called the morphogenetic furrow (MF) that sweeps anteriorly throughout the third instar larval stage and early pupal development. Anterior to the MF, cells undergo mitosis and are primed to differentiate. Posterior to the MF, each ommatidial column grows by sequential recruitment and differentiation of photoreceptors and non-neuronal cells. The R8 photoreceptor differentiates first, followed by the differentiation and recruitment of R2/5 and then R3/4 into the ommatidial cluster. All remaining undifferentiated cells then undergo another round of cell division known as the second mitotic wave (SMW). Photoreceptors R1/6, and R7 and non-neuronal cone cells differentiate from the pool of cells that exit the SMW and are recruited into growing ommatidial clusters. Pigment cells differentiate during pupal stages. A new ommatidial column forms every 2 hrs posterior to the MF such that each column is developmentally less mature than the column immediately posterior. Therefore, the larval eye disc is a spatiotemporal lattice with the most mature cells located at the posterior margin of the eye disc with less mature cells located anteriorly. This sequential arrangement of cells of different developmental ages is a unique aspect of the eye compared to most other *Drosophila* tissues. Cell type specification and differentiation from progenitors is highly regulated in the larval eye disc and has been extensively studied. However, much remains to be understood about early cell fate specification, determination, and differentiation. Although genome-wide genetic resources of the *Drosophila* eye (Michaut, Flister et al. 2003, Quiquand, Rimesso et al. 2021, Bollepogu Raja, Yeung et al. 2024), including single cell resources on the late larval and adult eye, have been reported (Yeung, Bollepogu Raja et al. 2022, Bollepogu Raja, Yeung et al. 2023), single cell RNA sequence (scRNA-seq) data from high quality cells that capture the cellular dynamics from the initiation of the MF to the onset of differentiation of each cell type in the larval eye is lacking.

In this study, we present single cell transcriptomic data generated from developing *Drosophila* larval eye discs from the initiation of the MF in late second instar larvae to the early stages of R1/6, R7 and cone cell differentiation in mid-larval eye discs, thus complementing previously published scRNA-seq studies on the *Drosophila* eye. Our data show cell clusters that correspond to all known major cell types present in early and mid-larval eye discs. We identify several cell type-specific markers and validate several such markers *in vivo*. Our data provide a valuable resource for investigating early stages of cell type differentiation in *Drosophila* and for researchers who use the larval eye disc as a model system.

## Results

### Single cell RNA sequencing of the developing *Drosophila* larval eye discs

We performed single cell RNA sequencing (scRNA-seq) from six time points during early larval *Drosophila* eye disc development using the 10x Genomics Chromium platform. To capture the gene expression dynamics from the initiation of the morphogenetic furrow (MF) to the differentiation of most major cell types, we first profiled eye disc cells from four time points spanning late second instar (69 hr after egg laying (AEL)) to third instar larval stages (72 hr, 75 hr and 78 hr AEL) (Figure 1A). The majority of eye disc cells at these time points are undifferentiated and actively dividing and only a minority of differentiated cell types are present. We therefore combined the data from these time points (termed “early larval”) for further analyses. Removal of dead or dying cells and multiplets, followed by dimension reduction using UMAP resulted in 3,624 eye cells. The UMAP plot shows clusters that correspond to all expected cell identities in the early larval eye disc (Figure 1B). As expected, the anterior undifferentiated (AUnd) cell cluster comprises the most abundant cell type (1723 cells) in the combined early larval data set. The AUnd cluster was identified using *Optix* (Seimiya and Gehring 2000), *eyeless* (*ey*) (Halder, Callaerts et al. 1998) and *teashirt* (*tsh*) (Pan and Rubin 1998), which are predominantly expressed in AUnd cells in developing larval eye discs (Figure 1C and 2B). The morphogenetic furrow (MF) cell cluster was identified using *decapentaplegic* (*dpp*) (Chanut and Heberlein 1997) and *rotund* (*rn*) (St Pierre, Galindo et al. 2002). These genes are expressed in the MF and *dpp* is required for the initiation and propagation of the MF across the eye disc (Chanut and Heberlein 1997) (Figure 1C and 2C,D). We also identified two cell populations that express known photoreceptor markers. One of these cell clusters is identified as R8, as it expresses *atonal* (*ato*) (Jarman, Grell et al. 1994, Baker, Yu et al. 1996) and *senseless* (*sens*) (Frankfort, Nolo et al. 2001). *ato* and *sens* are specifically expressed in R8 and are required for the differentiation of R8. *ato* is also expressed in the MF and in a stripe immediately anterior to the MF (Figure 1C and 2E,F). The second cell cluster expresses *rough* (*ro*) and *seven up* (*svp*) (Figure 1C and 2G,H). While *ro* is expressed in R2/5 and R3/4 (Kimmel, Heberlein et al. 1990), *svp* expression is confined to R3/4 and R1/6 (Mlodzik, Hiromi et al. 1990). Since this second cluster shows expression of both genes without segregation into individual photoreceptor subtype clusters, we reasoned that this cluster comprises the R2/5, R3/4 and R1/6 subtypes and therefore named this the R cell cluster. Interestingly, the AUnd, MF and arrangement of photoreceptor clusters in the cluster plot appears as a progression that is reminiscent of the temporal nature of the larval eye disc *in vivo*. The posterior undifferentiated cell cluster (PUnd) was identified using *Bar*-*H1* (*B-H1*) and *lozenge* (*lz*) which are known to be expressed in PUnd cells (Higashijima, Kojima et al. 1992, Flores, Daga et al. 1998) (Figure 1C and 2L). Our data also shows cell clusters corresponding to the peripodium (PPD) and the posterior cuboidal epithelium (PC). *Ance* and *ocelliless* (*oc*) mark the PPD (Bollepogu Raja, Yeung et al. 2023) (Figure 1C), while the pair rule genes *odd skipped* (*odd*), *drumstick* (*drm*) and *sister of odd and bowl* (*sob*) are confined to the PC (Figure 1C) and are required to initiate retinogenesis (Bras-Pereira, Bessa et al. 2006).

**Figure 1.**
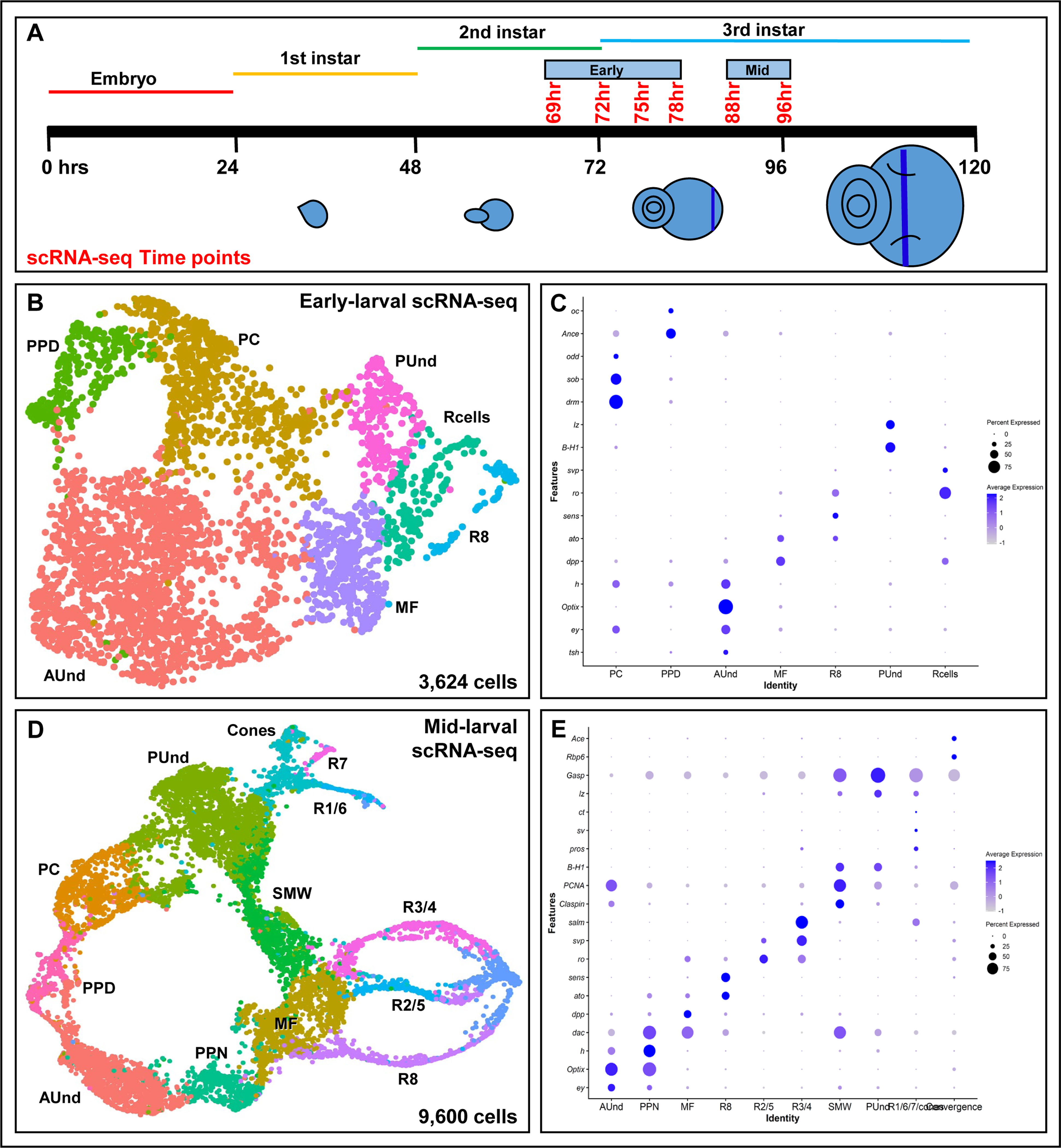
**A** Schematic of single cell RNA sequencing data generation from early to mid-larval eye discs. The time points reported in this work are shown in red. **B** scRNA-seq cluster plot generated from 3,624 early larval eye disc cells (69, 72, 75 and 78 hr after egg laying (AEL)) shows all expected cell identities. **C** DotPlot showing the expression of known markers. The size of the dot indicates the percentage of cells in a cluster showing expression of a given gene, while the intensity of the blue color shows the average expression level across all cells within each cluster. **D** scRNA-seq cluster plot generated from 9,600 mid-larval (88 and 96 hr AEL) eye disc cells with clusters corresponding to expected cell identities. **E** Dot plot showing the expression of mid-larval marker genes of different cell types. AUnd: Anterior Undifferentiated; PPN: Preproneural; MF: Morphogenetic Furrow; SMW: Second Mitotic Wave; PUnd: Posterior Undifferentiated; PC: Posterior Cuboidal Margin Peripodium; and PPD: Anterior Peripodial.

**Figure 2.**
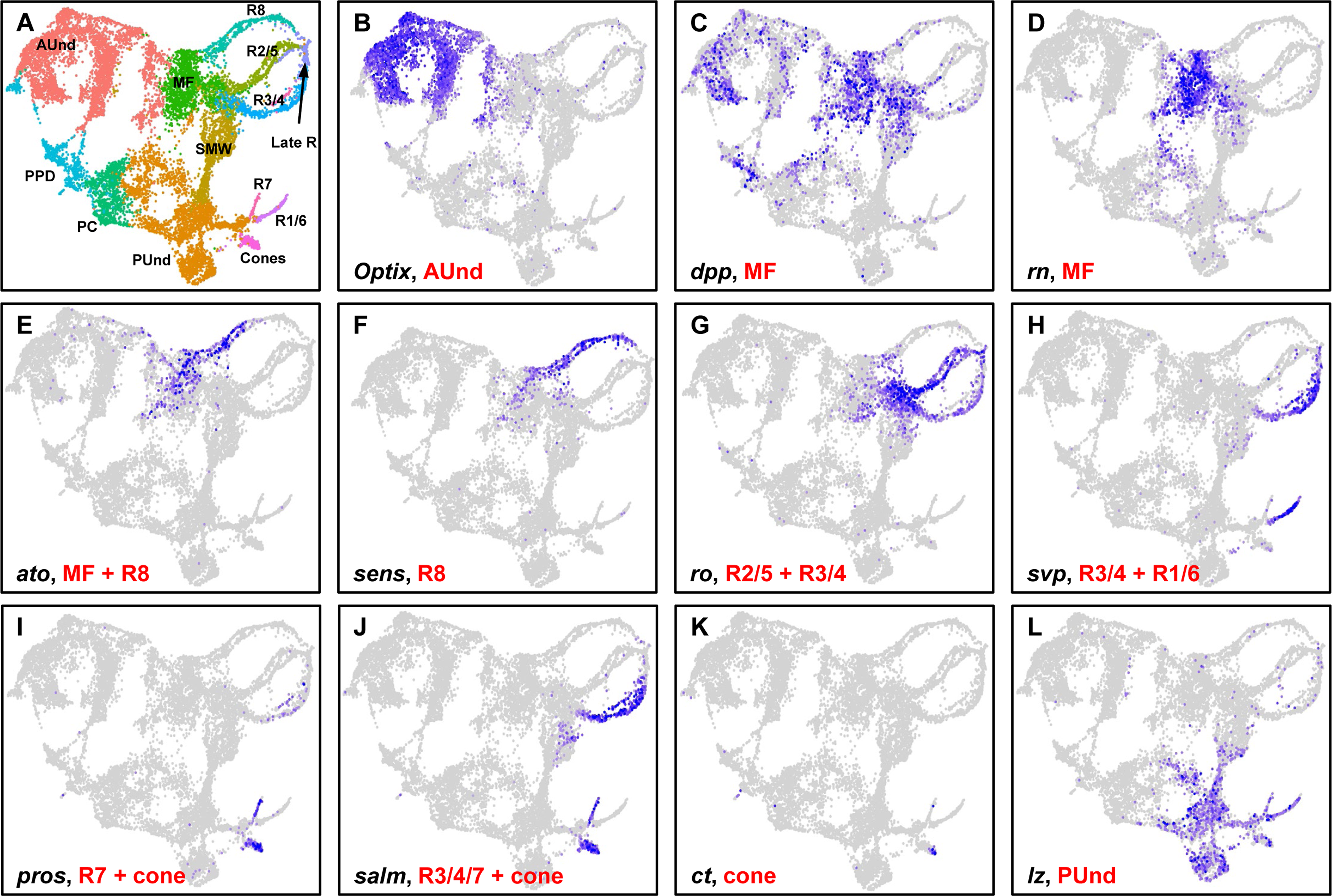
**A** scRNA-seq cluster plot generated by combining early and mid-larval data. **B**-**L** Feature Plots showing the expression (shown in blue) of selected marker genes that were used to identify and annotate clusters. The intensity of blue is proportional to the log-normalized expression levels. **B** *Optix* expression in AUnd cells. **C,D** *decapentaplegic* (*dpp*) and *rotund* (*rn*) expression in the MF. **E,F** *atonal* (*ato*) and *senseless* (*sens*) expression in the R8 cluster. *ato* is also expressed in the MF. **G** *rough* (*ro*) is expressed in the MF, R2/5, and R3/4. **H** *seven up* (*svp*) is expressed in R3/4 and R1/6. **I** *prospero* (*pros*) is expressed in R7 and cone cells. **J** *spalt major* (*salm*) is expressed in R3/4, R7 and cones. **K** *cut* (*ct*) is expressed in cones. **L** *lozenge* (*lz*) is expressed in PUnd cells and R1/6/7.

To complement our early larval scRNA-seq data, we performed scRNA-seq on two additional larval time points: 88 hr and 96 hr AEL. We combined these two data sets (termed “mid-larval”), filtered low quality cells and generated a UMAP cluster plot with 9,600 cells that shows all expected cell identities in the eye disc (Figure 1D). Similar to the early larval data, undifferentiated cell clusters were identified using *Optix* (AUnd), *hairy* (*h*) (Preproneural (PPN)), *dpp* (MF), *PCNA* (second mitotic wave (SMW)), and *lz* (PUnd) (Figure 1E). Photoreceptor cell clusters were identified using *sens* (R8), *ro* (R2/5 and R3/4), *svp* (R3/4 and R1/6), and *prospero* (*pros*) (Kauffmann, Li et al. 1996) (R7) (Figure 1E and 2I,K). Cone cells were identified using *cut* (*ct*) (Blochlinger, Jan et al. 1993) and *pros* expression. *pros* is expressed in both R7 and cone cells (Figure 1E and 2I). Importantly, the mid-larval eye scRNA-seq cluster plot closely resembles our previously published late larval cluster plot (Bollepogu Raja, Yeung et al. 2023) with each cell cluster appearing with progressive patterns of gene expression suggesting a temporal component, reminiscent of the temporal nature of the larval eye disc. The photoreceptor clusters appear as strands of cells that are either connected to the MF (R8, R2/5 and R3/4) or the PUnd (R1/6 and R7) cell clusters. Though our mid-larval cluster plot shows R1/6, R7 and cone cells, they do not segregate distinctly and appear together as one cluster. Taken together, our early and mid-larval data show that we captured all major types from eye discs.

### Gene expression profiles of anterior undifferentiated cells

Since our early larval data profiles only ∼3,600 cells, we combined early larval and mid-larval data for downstream analyses. This merging of data enables good representation of each cell identity as well as profiles cell types at high resolution. This is possible because each larval eye disc is a spatiotemporal continuum with cells of different developmental ages in the same tissue. The combined data showed all expected cell identities that were observed in both early and mid-larval time points (Figure 2A). We then performed differential gene expression analyses between cell clusters to understand their gene expression profiles. As expected, *Optix*, *tsh*, *twin of eyeless* (*toy*) and *ey* are among the top 15 differentially expressed markers in the AUnd cell cluster (Supplemental Data 1). It is well known that these genes are expressed in undifferentiated cells prior to the initiation of the MF as well as anterior to the MF after initiation (Halder, Callaerts et al. 1998, Pan and Rubin 1998, Seimiya and Gehring 2000). We also identified *CG17211* and *Organic anion transporting polypeptide 74D* (*Oatp74D*) (Figure 3B,C) as two examples of novel AUnd markers. These are expressed predominantly in the AUnd cell cluster in which *Optix* is expressed and are likely AUnd markers. The complete list of AUnd markers is shown in Supplemental Data 1. We further performed Gene Ontology (GO) term enrichment using these marker genes and, as expected, terms related to imaginal disc development and proliferation showed enrichment (Supplemental Data 2). These include eye-antennal disc development (GO:0035214) and positive regulation of cell population proliferation (GO:0008284).

**Figure 3.**
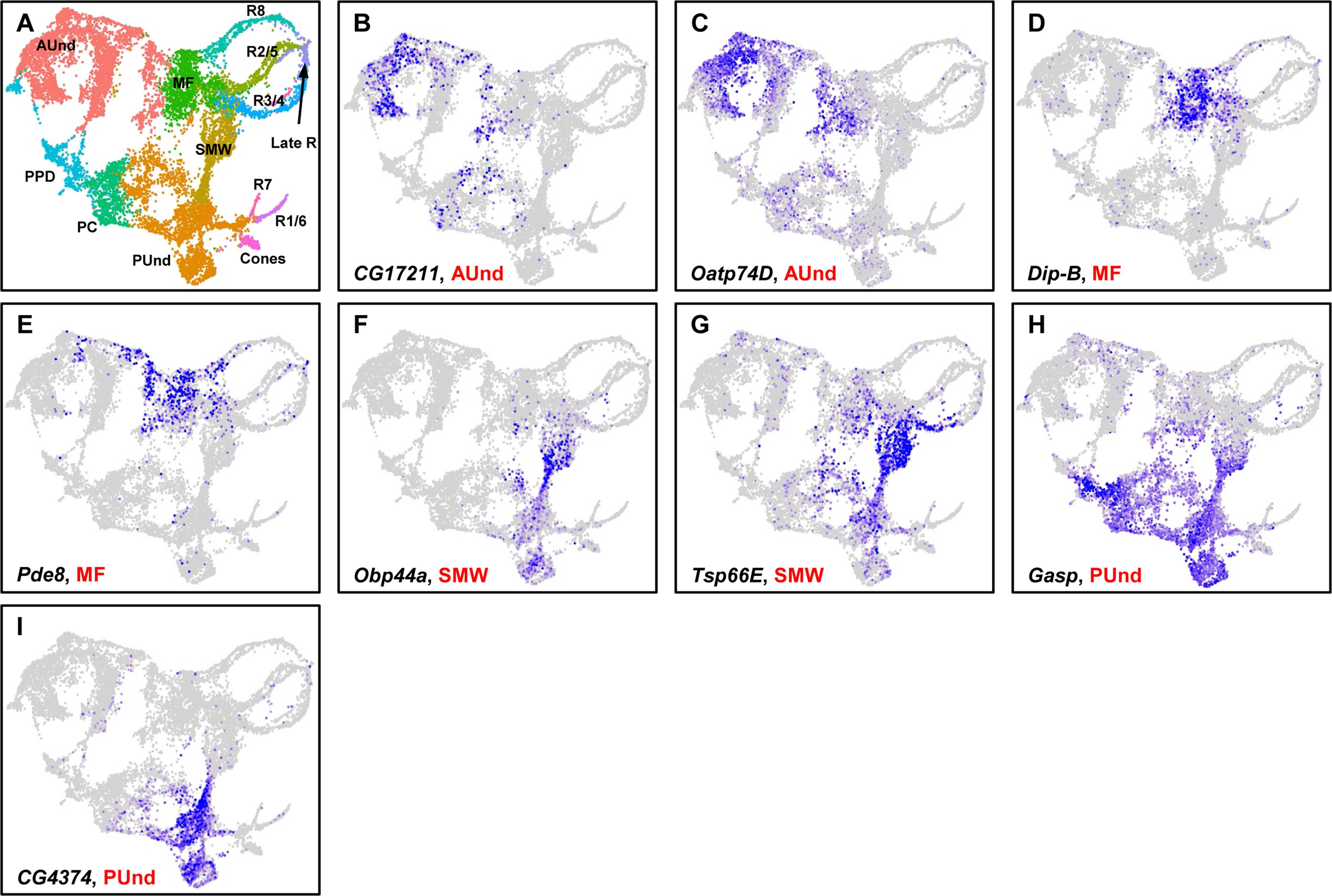
Markers of undifferentiated cells. **A** UMAP cluster plot of the combined early and mid-larval eye disc data. **B**-**I** Feature Plots showing the expression of cell type-specific genes. *CG17211* (**B**) and *Oatp74D* (**C**) are expressed in the AUnd. *Dip-B* (**D**) and *Pde8* (**E**) mRNA is detected in the MF. *Obp44a* (**F**) and *Tsp66E* (**G**) are predominantly expressed in the SMW cluster. *Gasp* (**H**) and *CG4374* (**I**) expression are observed in the PUnd.

We also performed principal component analysis (PCA) on the AUnd cell cluster to identify genes that change as a function of time and may be involved in dynamic cellular processes such as cell division and differentiation. As expected, PCA on the AUnd cluster identified *Optix* with the second highest loading weight. The long non-coding RNA *lncRNA:CR33938* is the top PC1 gene for the PUnd cluster. *Odorant-binding protein 56a* (*Obp56a*), *Oatp74D*, fatty acid binding protein (*fabp*) and *narrow* (*nw*) are the other top 6 PC1 genes. The top 1000 PC1 genes of the AUnd cluster with loading weights are shown in Supplemental Data 3. These top PC1 genes may be involved in dynamic processes of the AUnd cell cluster such as cell division, growth, and differentiation.

### Gene expression profiles of the morphogenetic furrow

Differential gene expression analyses of the MF cluster show five *Enhancer of split* (*E(spl)*) genes in the top 10 marker gene list (Supplemental Data 1). *E(spl)* genes are the downstream effectors of Notch signaling pathway (Cooper and Bray 1999). The role of Notch signaling in the initiation and propagation of MF is well known (Cooper and Bray 1999) and it is expected that the *E(spl)* genes appear as differentially expressed genes in the MF. In addition to other known MF genes such as *dpp* and *rn*, *Dipeptidase B* (*Dip-b*) and *Phosphodiesterase 8* (*Pde8*) (Figure 3D,E) are predominantly expressed in MF and may play a role in MF initiation and/or progression.

To identify putative maturation genes, we subjected the MF cluster to PCA. As expected, the *E(spl)* genes show the highest loading weights in PC1 along with other known MF genes such as *rn*. Similarly, *scabrous* (*sca*) and *ato* are in the top 30 PC1 genes. These genes are involved in the specification of R8 from the pool of undifferentiated progenitor cells (Jarman, Grell et al. 1994, Lee, Hu et al. 1996). In addition, *CG15282*, *gliolectin* (*glec*), *Protein kinase cAMP dependent regulatory subunit type 2* (*Pka-R2*), *Bearded* (*Brd*) and *Dip-*B are other top 30 PC1 genes whose function in the progression and maturation of cells in MF has not been previously reported. Taken together, these data suggest that we have identified several potential candidate genes that may drive specification of cell types in the MF.

### Cell cycle and DNA replication genes are enriched in the second mitotic wave cluster

Differential gene expression analyses of the SMW cluster reveals known markers such as *dac* and the cell cycle-related genes *Claspin* and *PCNA* (Bollepogu Raja, Yeung et al. 2023). In addition, *Odorant-binding protein 44a* (*Obp44a*) and *Tetraspanin 66E* (*Tsp66E*) (Figure 3F,G) and other markers were also identified and are shown in Supplemental Data 1. GO term analyses shows enrichment for terms involved in the cell cycle and DNA replication. Some of the GO terms include pyrimidine deoxyribonucleoside monophosphate biosynthetic process (GO:0009177), deoxyribonucleotide biosynthetic process (GO:0009263), leading strand elongation (GO:0006272) and cell cycle DNA replication initiation (GO:1902292). In particular, the DNA replication licensing factor and Minichromosome maintenance (MCM) complex genes (Su and O’Farrell 1997, Su and O’Farrell 1998), which are replicative helicases for DNA replication, initiation and elongation, are associated with these GO terms. This is expected as the cells in the SMW are actively undergoing cell division and preparing to differentiate into R1/6, R7 or cone cells. We then subjected the SMW cluster to PCA to identify putative genes that may prime these cells to undergo division and differentiation. *anterior open* (*aop*/*yan*) shows the highest loading weight in PC1 of the SMW. The role of *aop* in regulating cell fate transitions in the eye is well documented (O’Neill, Rebay et al. 1994). *target of wit* (*twit*), *Obp44a*, *CG42342* and *Tsp66E* are the other top 5 PC1 genes in the SMW, whose function in the SMW is currently unknown.

### Gene expression profiles of posterior undifferentiated cells

Our cluster plot shows that the PUnd cells cluster separately from AUnd cells, reflecting transcriptomic differences between undifferentiated cells anterior and posterior to the MF. PUnd cells are developmentally more mature progenitors compared to AUnd cells and it is expected that the two undifferentiated cell populations will segregate on the UMAP plot (Bollepogu Raja, Yeung et al. 2023). In addition to the known PUnd markers *lz* and *B-H1*, differential gene expression analyses identify several PUnd markers such as *Gasp* and *CG4374* (Figure 3H,I). These genes are specifically expressed in the PUnd cell cluster and their function is currently not known. PCA on PUnd cell cluster identified *CG13071*, *CG9691*, *Gasp*, *kekkon 1* (*kek1*) and *scarface* (*scaf*) as the top 5 genes with the most variation in the PUnd cluster. GO analyses of the PUnd cell cluster showed enrichment for negative regulation of compound eye photoreceptor cell differentiation (GO:0110118), positive regulation of compound eye retinal cell programmed cell death (GO:0046672) and negative regulation of cell fate specification (GO:0009996). These terms are expected because photoreceptors have undergone differentiation and some of the undifferentiated cells are eliminated through apoptosis.

### Photoreceptor subtypes and cones cells appear as distinct clusters

Our data show five cell populations that appear as distinct strands, which we identified as photoreceptor subtypes (Figure 2A). Three of these strands are connected to the MF and correspond to R8, R2/5 and R3/4, whereas R1/6 and R7 are connected to the PUnd cell cluster. This arrangement of clusters resembles the sequence of differentiation of cell types in the physical eye disc. R8, R2/5 and R3/4 differentiate first from cells in the MF, while R1/6, R7 and cones differentiate after the SMW and therefore are derived from posterior undifferentiated cells. Similar to our late larval data set (Bollepogu Raja, Yeung et al. 2023), photoreceptor cells cluster as temporal strands with less mature cells located near the MF or PUnd clusters, while more mature cells are located at opposite ends of each strand. The expression of *sens*, *ato* and *bride of sevenless* (*boss*) reveals the temporal nature of the R8 strand (Figure 4A-C). *ato* is expressed in the MF and in less mature R8 cells, *sens* is expressed in all R8 cells, and *boss* is expressed only in mature R8 cells. Our data show *ato* expression in the MF cluster and proximal tip of the R8 strand that is close to the MF but no *ato* expression is observed in the rest of the strand (Figure 4A). *sens* mRNA is detected along the entire stream (Figure 4B), while *boss* expression is observed only in mature R8 cells that are at the distal tip of the R8 strand (Figure 4C). These data collectively indicate that the R8 photoreceptor strand is a continuum of cells arranged progressively according to their developmental age. Likewise, other photoreceptor subtype streams also exhibit such temporal dynamics. In addition, similar to our late larval eye scRNA-seq data, the R8, R2/5 and R3/4 strands appear to connect to a common cluster at their more mature ends and is named the ’Late R’ cell cluster (Late R, Figure 3A) (Bollepogu Raja, Yeung et al. 2023). Genes related to axon projection, guidance and synapse formation dominate the differentially expressed gene list of the Late R cell cluster. Cone cells appear as a separate cluster that is connected to the PUnd cluster. Our cluster plot shows that R1/6, R7 and cones are distinct but are part of one major cluster. Differential gene expression analyses identified many photoreceptor and cone subtype-specific markers and these are shown in Supplemental Data 1.

**Figure 4.**
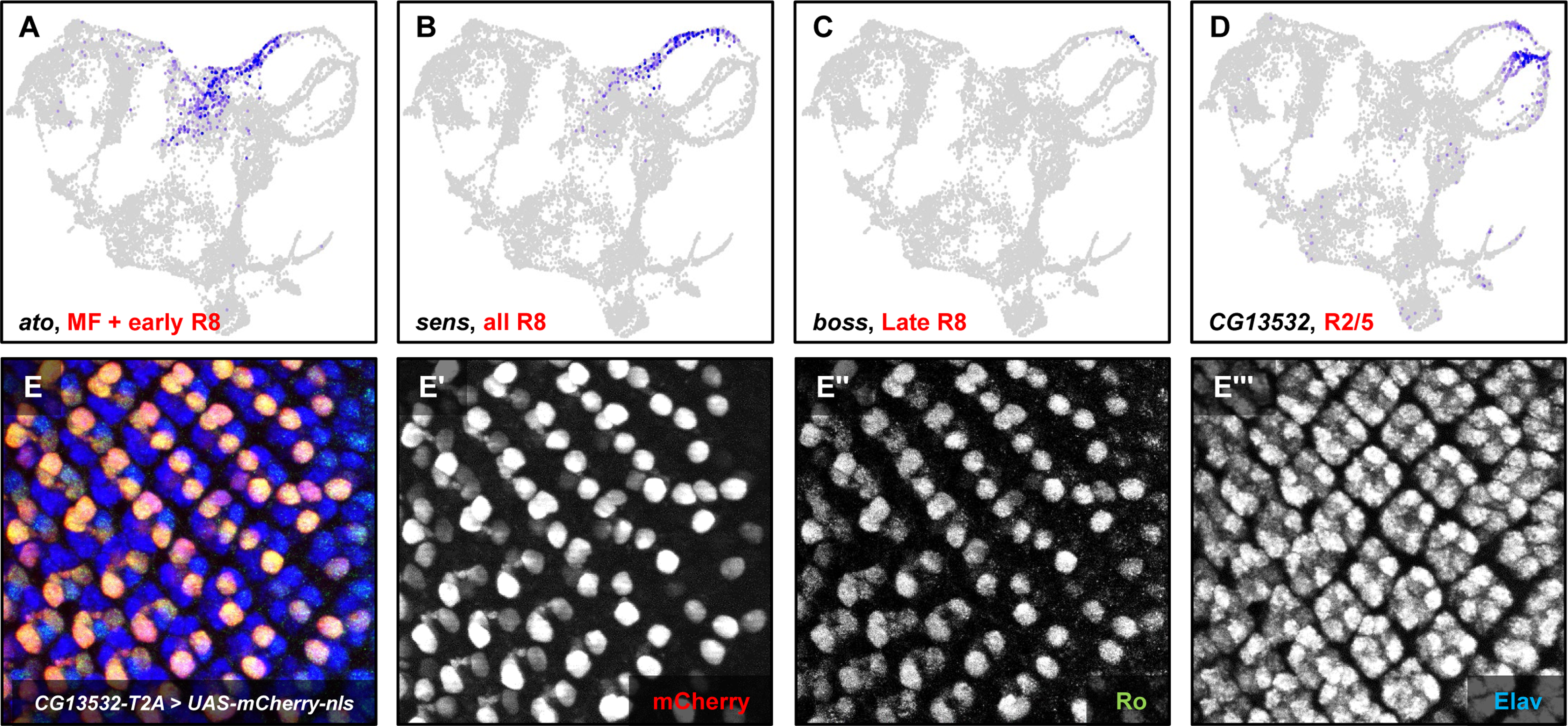
Distinct strands of photoreceptor cells exhibit a spatiotemporal dimension. **A** Feature Plot showing the expression of *ato* in the MF and R8 near the MF. **B** Feature Plot showing *sens* expression distributed throughout the R8 strand. **C** Feature Plot showing *boss* expression in more mature R8 cells (far, upper right of the plot). **D** *CG13532* is predominantly expressed in R2/5 and is also observed at lower levels in mature R8 and R3/4 cells. **E**-**E’’’** Staining of eye discs from *CG13532-T2A-Gal4*□>□*UAS-nls-mCherry* larvae showing mCherry expression in Ro-positive cells.

As an *in vivo* validation step we used *Trojan-Gal4* (*T2A-Gal4*) lines (Lee, Zirin et al. 2018) to test the expression of marker genes in the larval eye disc. *T2A-Gal4* insertions express Gal4 under the control of endogenous promoters and recapitulate the endogenous pattern of expression of the gene in which the transgene is inserted. Our scRNA-seq data show that *CG13532* is predominantly expressed in R2/5 and some expression is also observed in late R8. We used *CG13532-T2A-Gal4* to drive expression of a nuclear localized mCherry reporter (*UAS-mCherry-nls*) and costained larval eye discs with mCherry, Rough (Ro, an R2/5 marker) and Embryonic lethal abnormal vision (Elav, a pan-neuronal marker) antibodies. We observed that mCherry, Ro and Elav colocalize in two cells per ommatidium posterior to MF (Figure 4E-E’’’) suggesting that *CG13532* is expressed in R2/5 cells and is an R2/5 marker. Furthermore, these results suggest that our scRNA-seq data can predict the expression and distribution of unknown genes in the larval eye disc.

### Gene expression profiles of peripodial cells are distinct from other cell types

The peripodial membrane is a thin squamous epithelium that covers the eye disc proper and is important for its patterning and growth. In addition, short narrow cells comprising the ’posterior cuboidal margin’ (PC) are also present at the posterior border between the eye disc and peripodial layers. Our data show two clusters that correspond to the peripodial cells (PPD) and the PC (Figure 3A). The PPD was identified using *Ance* and *oc*, which are expressed in the PPD. Differential gene expression analyses of the PPD cluster identified several PPD markers including *Odorant-binding protein 99a* (*Obp99a*) and *CG14984*, which are expressed in the PPD (Figure 5A,B). *CG14984* is also expression in the AUnd. Several other larval eye PPD markers are shown in Supplemental Data 1. As expected, GO term enrichment analyses of the PPD shows terms related to tissue morphogenesis and appendage development as some of the head structures are derived from cells of the PPD. Examples of these terms include muscle tissue morphogenesis (GO:0060415), imaginal disc-derived appendage morphogenesis (GO:0035114), and imaginal disc development (GO:0007444). Surprisingly, we also see axon-related terms in our GO analyses such as axon guidance (GO:0007411), neuron projection guidance (GO:0097485) and neuron projection morphogenesis (GO:0048812).

The PC cluster shows expression of the pair-rule genes *sob*, *odd* and *drm*, which are specifically expressed in this cluster and trigger retinogenesis in the eye disc along with *dpp* and *dac* (Bras-Pereira, Bessa et al. 2006). *dac* expression is also observed in this cluster. *dac* is required for proper cell fate determination in the eye imaginal disc and cells at the posterior margin of the eye disc adopt a cuticle fate in the absence of *dac* function (Pappu, Ostrin et al. 2005). Differential gene expression analyses identified *CG17278* and *crossveinless c* (*cv-c*) (Supplemental Figure 1C,D) as PC markers that are specifically expressed in this cluster. The function of these genes in PC cells and how they affect eye development is currently unknown. We also performed GO analyses using PC differentially expressed genes and, interestingly, terms related to plasma membrane extension show enrichment. These include cytoneme assembly (GO:0035231), cytoneme morphogenesis (GO:0003399) and filopodium assembly (GO:0046847). In addition, chitin-based cuticle attachment to epithelium (GO:0040005) and larval chitin-based cuticle development (GO:0008363) GO terms are also associated with genes in the PC cluster. This is expected as peripodial cells are known to form adult head cuticle (Stultz, Lee et al. 2006).

## Discussion

Here we report a single cell RNA-seq atlas of the developing *Drosophila* larval eye disc at early and mid-larval time points. We profiled a total of ∼13,000 cells, which is several fold more than the number of cells present in a single disc averaged over these time points. Since there are fewer subtypes of cells in this tissue prior to differentiation, each type should be well represented in our dataset. Both our early-and mid-larval data show distinct cell clusters corresponding to all major known cell identities present in the eye disc at these time points. To mitigate stress-induced transcription, we performed dissection and dissociation of eye discs in the presence of the transcription inhibitor Actinomycin-D (ActD) (Yeung, Bollepogu Raja et al. 2022, Bollepogu Raja, Yeung et al. 2023). Therefore, the gene expression patterns observed in our study most likely accurately reflect the endogenous mRNA patterns. Analyses of our data reveal many putative markers for each cell type, several of which have been validated using *T2A-Gal4* drivers *in vivo*. The data generated in this study will likely benefit research groups that use the larval eye disc as a model system and compliments our previously published late larval and adult eye scRNA-seq data (Yeung, Bollepogu Raja et al. 2022, Bollepogu Raja, Yeung et al. 2023).

A majority of cells in early larval eye discs are undifferentiated progenitor cells that are either actively dividing or are primed to undergo differentiation into different cell types. Our data show clusters corresponding to several undifferentiated cell populations (AUnd, MF, SMW, PUnd) with distinct transcriptomic profiles. Further, our analyses have revealed many putative markers for all undifferentiated cell types. Investigating the transcriptomic profiles of undifferentiated cell clusters may reveal mechanisms that underlie specification, differentiation and survival of cell types. For instance, our data show many markers in the AUnd, MF and PUnd cell clusters and the function of most of these genes in *Drosophila* eye development is unknown. Gene network analyses and functional testing of these genes may unravel their significance in driving the specification and differentiation of distinct cell types in the eye disc. Similarly, the SMW cell cluster expresses genes that regulate the cell cycle, initiation of DNA replication and repair, and chromatin remodeling, as inferred from GO enrichment analyses. These gene lists can be a valuable resource for groups that study different aspects of cell division and survival. Furthermore, since most *Drosophila* genes are highly conserved with humans, the data generated in this report will likely benefit research groups that use the larval eye disc as a model system to study genes that may underlie diseases processes in humans (Wawersik and Maas 2000, Sang and Jackson 2005, Prüßing, Voigt et al. 2013).

Our mid-larval data also show cell clusters that correspond to all major cell types present in the eye. Similar to our late larval scRNA-seq (Bollepogu Raja, Yeung et al. 2023), each photoreceptor cell cluster appears as a progression, with strands of cells that are connected to the MF or the PUnd. Each photoreceptor strand is a space-time continuum with cells arranged along the strand according to their developmental age. More mature cells are located at the distal tips of each strand, while the less mature cells are near the MF or PUnd clusters, similar to the position of photoreceptors in the physical eye disc. Since R1/6 and R7 have just begun to differentiate at the mid-pupal time point, we do not yet see fully extended strands of these subtypes as observed in the late larval scRNA-seq data. Interestingly, our mid-larval data also show the Late R cell cluster, which comprises mature photoreceptor cells that express axon projection genes. The data presented in this report, along with the published late larval data (Bollepogu Raja, Yeung et al. 2023), cover the full repertoire of gene expression of distinct cell types in the eye disc from late second instar larvae (69 hr APF) to late third instar developmental stages. Together, these data can be used as a resource to study how genes and pathways change as a function of time to drive development and maturation of different cell types.

Our data show two peripodial cell clusters, PPD and PC. One of these clusters comprise peripodial cells that cover the entire larval eye disc (PPD), while the other cluster consists of cuboidal cells located at the posterior margin of the eye disc (PC). The peripodial membrane is required for eye pattern formation and coordinates growth as well as patterning of the eye disc proper (Cho, Chern et al. 2000, Gibson and Schubiger 2000). The morphogens *wingless* (*wg*), *dpp* and *hedgehog* (*hh*) are expressed in the peripodial membrane and are required for proper growth (Cho, Chern et al. 2000). Loss of *hh* in peripodial cells disrupts growth in the eye disc proper. In addition, retinogenesis is induced from the posterior cuboidal epithelium by the *odd skipped* gene family members *odd*, *drm* and *sob*, which are specifically expressed in the PC (Bras-Pereira, Bessa et al. 2006). Peripodial cells, therefore, are crucial for development and pattern formation in the eye. These functions and signaling between peripodial cells and the eye disc proper may be mediated by cellular processes, called axonemes, that extend into the eye disc proper (Cho, Chern et al. 2000). However, not much is known about how these processes develop or the genes that produce these cellular extensions. Our data show markers that are expressed in the PPD and PC and the function of most of these genes in eye development has not been reported. In addition, GO analyses using differentially expressed genes in peripodial cell clusters show enrichment for terms associated with tracheal growth and tracheal lumen proteins. These genes may be involved in the formation of translumenal extensions between peripodial and eye cells and provide new avenues for future research on cell signaling and cell communication.

In summary, we report single cell transcriptomic atlases of the *Drosophila* larval eye from the initiation of MF progression to the mid-larval time stage. All major cell types present in the developing larval eye discs are represented in these data. We identified many cell type markers and validated several of them *in vivo*. The function of most of these genes in *Drosophila* eye development is unclear and therefore these single cell resources expand the wealth of genome-wide transcriptomic data that is available on the larval eye and will aid research involving cell fate determination, development, and function.

## Materials and Methods

### Fly husbandry

All flies used in this study were maintained at 25°C on cornmeal agar medium. We used larval eye discs from the *Drosophila melanogaster Canton-S* strain for all single cell experiments. The following fly stocks were obtained from the Bloomington *Drosophila* Stock Center: *UAS-mCherry-nls* (38424), *CG13532* (91332).

### Dissociation protocol to generate single cells from larval eye discs

30 to 40 eye discs were dissected in 1x PBS supplemented with 1.9 µM Actinomycin-D (a known transcription inhibitor) and were immediately transferred to a LoBind 1.5 ml Eppendorf tube containing 700 µl ice-cold Rinaldini insect solution and 1.9 µM Actinomycin-D. Early larval eye discs were dissociated by adding 16 µl of collagenase (100 mg/ml; Sigma-Aldrich #C9697), while mid-larval eye discs were dissociated with 16 µl collagenase and 2 µl of dispase (1mg/ml; Sigma-Aldrich #D4818). The tube was then placed horizontally in a shaker and eye discs were dissociated at 32°C at 250 rpm. The solution was pipetted every 10 min to disrupt clumps of cells and the extent of dissociation was examined every 10 min. After 45 min, the dissociation was stopped by adding 1 ml of Rinaldini solution containing 0.05% Bovine Serum Albumin (BSA). The cell suspension was filtered using a 35 µm sterile filter and centrifuged at 4°C at 50 to 100 rcf to obtain a cell pellet. The pellet was washed once with Rinaldini + 0.05% BSA, gently mixed and was subjected to centrifugation. The cell pellet was gently resuspended in Rinaldini + 0.05% BSA using a wide-bore pipette tip. Viability was assessed using Hoechst-propidium iodide and cell suspensions with >95% viability were used for scRNA-seq experiments at a concentration of 1000 to 1200 cells/µl.

### Single cell RNA-seq using 10x Genomics

Single cell libraries were generated using the Chromium Next GEM Single Cell 3’ Reagent Kit v3.1 from 10x Genomics. Briefly, single cell suspensions were mixed with Gel Beads containing barcodes and loaded on a 10x Genomics Chromium Controller which isolates each cell in an oil droplet with a Gel Bead (GEM). The cells are lysed within the bead, the mRNA is captured and barcoded before synthesizing cDNA by reverse transcription. A library was generated from pooled cDNAs and sequenced with a NovaSeq 6000 (Illumina). FASTQ files generated from each sequencing run were initially analyzed using the Cell Ranger v6.0.1 count pipeline. The reference genome was built using *Drosophila melanogaster* reference genome Release 6 (dm6).

### Seurat Analyses

The filtered gene expression matrices from the Cell Ranger output were used as input to perform downstream analyses in Seurat v4.03. We first removed potential multiplets and lysed cells by removing cells that showed a total number of genes below 200 or above 4000 and cells that showed high mitochondrial gene percentage (>30%). The remaining cells were then normalized and the data was scaled across all cells using the SCTransform algorithm. The total number of variable features used for SCTransform was 4000. We then performed regression using mitochondrial genes and the data was then reduced using the top 50 dimensions. The data was then clustered using the RunUMAP, FindNeighbors and FindClusters functions in Seurat. Eye disc, PPD and PC cells were retained and non-retinal cells were removed using known markers. These include antenna (*Distal-less* (*dll*)) (Dong, Dicks et al. 2002), glia (*reversed polarity* (*repo*)) (Xiong, Okano et al. 1994) and brain (*found in neurons* (*fne*)) (Samson and Chalvet 2003). Differential marker gene lists for all cell clusters were generated using a log-fold change threshold value of 0.25 and a minimum percentage of cells in which the gene is detected of 25%. The Seurat merge function was used to combine data from different time points and SeuratWrapper function RunHarmony was used to remove batch effects.

### Immunohistochemistry

Larval eye discs were dissected and fixed in 3.7% paraformaldehyde in PBS for 30 min at room temperature. Eye discs were then washed 3 times with PBT (PBS+0.3% Triton X-100) and blocked using 5% normal goat serum in PBT. Primary antibody incubations were done overnight at 4°C. Secondary antibody incubations were performed at room temperature for at least 1 hr. Optically stacked images were generated using a Zeiss Apotome Imager microscope. The images were processed with Zen Blue and Adobe Photoshop software. We used the following antibodies: rat anti-Elav (DHSB-7E8A10, RRID:AB, #52818, 1:500), rabbit anti-mCherry (Thermo Fischer Scientific, catalog number: MA5-47061,RRID:AB, #2889995, 1:2000) and mouse anti-Rough (DHSB-62C2A8, RRID: AB, #528456, 1:500).

### GO term enrichment analyses

Differentially expressed genes from each cell cluster were used for analysis with Panther (Mi, Lazareva-Ulitsky et al. 2005) using default values.

---

**Supplemental Figure 1.**
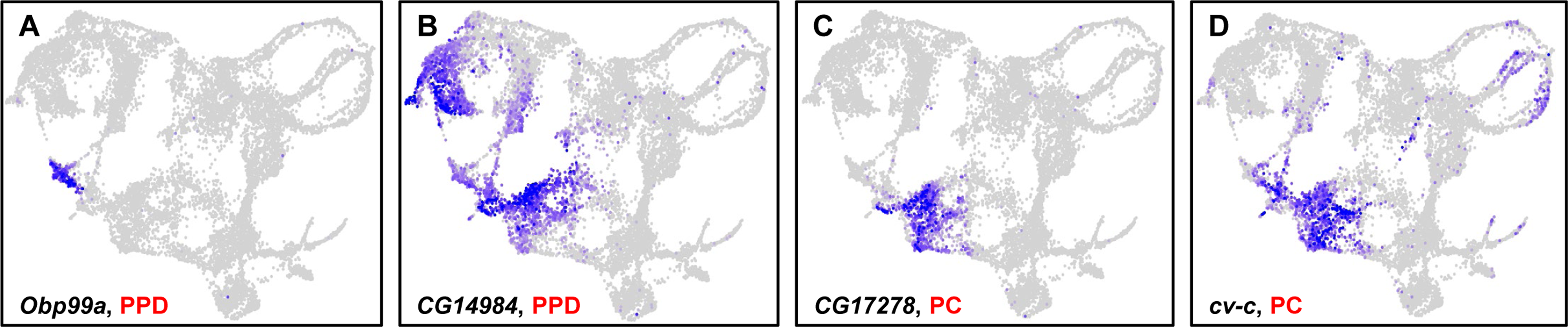

## References

Baker, N. E., S. Yu and D. Han (1996). “Evolution of proneural atonal expression during distinct regulatory phases in the developing Drosophila eye.” Current Biology 6(10): 1290–1302.

Baslan, T. and J. Hicks (2017). “Unravelling biology and shifting paradigms in cancer with single-cell sequencing.” Nature Reviews Cancer 17(9): 557–569.

Blochlinger, K., L. Y. Jan and Y. N. Jan (1993). “Postembryonic patterns of expression of cut, a locus regulating sensory organ identity in Drosophila.” Development 117(2): 441–450.

Bollepogu Raja, K. K., K. Yeung, Y.-K. Shim, Y. Li, R. Chen and G. Mardon (2023). “A single cell genomics atlas of the Drosophila larval eye reveals distinct photoreceptor developmental timelines.” Nature Communications 14(1): 7205.

Bollepogu Raja, K. K., K. Yeung, Y.-K. Shim and G. Mardon (2024). “Integrative genomic analyses reveal putative cell type-specific targets of the Drosophila ets transcription factor Pointed.” BMC Genomics 25(1): 103.

Bras-Pereira, C., J. Bessa and F. Casares (2006). “Odd-skipped genes specify the signaling center that triggers retinogenesis in Drosophila.”

Chanut, F. and U. Heberlein (1997). “Role of decapentaplegic in initiation and progression of the morphogenetic furrow in the developing Drosophila retina.” Development 124(2): 559–567.

Cho, K.-O., J. Chern, S. Izaddoost and K.-W. Choi (2000). “Novel signaling from the peripodial membrane is essential for eye disc patterning in Drosophila.” Cell 103(2): 331–342.

Consortium, T. M. (2018). “Single-cell transcriptomics of 20 mouse organs creates a Tabula Muris.” Nature 562(7727): 367–372.

Cooper, M. T. and S. J. Bray (1999). “Frizzled regulation of Notch signalling polarizes cell fate in the Drosophila eye.” Nature 397(6719): 526–530.

Davie, K., J. Janssens, D. Koldere, M. De Waegeneer, U. Pech, Ł. Kreft, S. Aibar, S. Makhzami, V. Christiaens and C. B. González-Blas (2018). “A single-cell transcriptome atlas of the aging Drosophila brain.” Cell 174(4): 982–998. e920.

Dong, P. S., J. S. Dicks and G. Panganiban (2002). “Distal-less and homothorax regulate multiple targets to pattern the Drosophila antenna.”

Flores, G. V., A. Daga, H. R. Kalhor and U. Banerjee (1998). “Lozenge is expressed in pluripotent precursor cells and patterns multiple cell types in the Drosophila eye through the control of cell-specific transcription factors.” Development 125(18): 3681–3687.

Frankfort, B. J., R. Nolo, Z. Zhang, H. Bellen and G. Mardon (2001). “senseless repression of rough is required for R8 photoreceptor differentiation in the developing Drosophila eye.” Neuron 32(3): 403–414.

Gibson, M. C. and G. Schubiger (2000). “Peripodial cells regulate proliferation and patterning of Drosophila imaginal discs.” Cell 103(2): 343–350.

Halder, G., P. Callaerts, S. Flister, U. Walldorf, U. Kloter and W. J. Gehring (1998). “Eyeless initiates the expression of both sine oculis and eyes absent during Drosophila compound eye development.” Development 125(12): 2181–2191.

Higashijima, S.-i., T. Kojima, T. Michiue, S. Ishimaru, Y. Emori and K. Saigo (1992). “Dual Bar homeo box genes of Drosophila required in two photoreceptor cells, R1 and R6, and primary pigment cells for normal eye development.” Genes & Development 6(1): 50–60.

Jarman, A. P., E. H. Grell, L. Ackerman, L. Y. Jan and Y. N. Jan (1994). “Atonal is the proneural gene for Drosophila photoreceptors.” Nature 369(6479): 398–400.

Karaiskos, N., P. Wahle, J. Alles, A. Boltengagen, S. Ayoub, C. Kipar, C. Kocks, N. Rajewsky and R. P. Zinzen (2017). “The Drosophila embryo at single-cell transcriptome resolution.” Science 358(6360): 194–199.

Kauffmann, R. C., S. Li, P. A. Gallagher, J. Zhang and R. W. Carthew (1996). “Ras1 signaling and transcriptional competence in the R7 cell of Drosophila.” Genes & Development 10(17): 2167–2178.

Kimmel, B. E., U. Heberlein and G. M. Rubin (1990). “The homeo domain protein rough is expressed in a subset of cells in the developing Drosophila eye where it can specify photoreceptor cell subtype.” Genes & Development 4(5): 712–727.

Kolodziejczyk, A. A., J. K. Kim, V. Svensson, J. C. Marioni and S. A. Teichmann (2015). “The technology and biology of single-cell RNA sequencing.” Molecular cell 58(4): 610–620.

Lee, E.-C., X. Hu, S.-Y. Yu and N. E. Baker (1996). “The scabrous gene encodes a secreted glycoprotein dimer and regulates proneural development in Drosophila eyes.” Molecular and Cellular Biology.

Lee, P.-T., J. Zirin, O. Kanca, W.-W. Lin, K. L. Schulze, D. Li-Kroeger, R. Tao, C. Devereaux, Y. Hu and V. Chung (2018). “A gene-specific T2A-GAL4 library for Drosophila.” Elife 7.

Lim, B., Y. Lin and N. Navin (2020). “Advancing cancer research and medicine with single-cell genomics.” Cancer cell 37(4): 456–470.

Mi, H., B. Lazareva-Ulitsky, R. Loo, A. Kejariwal, J. Vandergriff, S. Rabkin, N. Guo, A. Muruganujan, O. Doremieux and M. J. Campbell (2005). “The PANTHER database of protein families, subfamilies, functions and pathways.” Nucleic acids research 33(suppl_1): D284-D288.

Michaut, L., S. Flister, M. Neeb, K. P. White, U. Certa and W. J. Gehring (2003). “Analysis of the eye developmental pathway in Drosophila using DNA microarrays.” Proceedings of the National Academy of Sciences 100(7): 4024–4029.

Mlodzik, M., Y. Hiromi, U. Weber, C. S. Goodman and G. M. Rubin (1990). “The Drosophila seven-up gene, a member of the steroid receptor gene superfamily, controls photoreceptor cell fates.” Cell 60(2): 211–224.

O’Neill, E. M., I. Rebay, R. Tjian and G. M. Rubin (1994). “The activities of two Ets-related transcription factors required for Drosophila eye development are modulated by the Ras/MAPK pathway.” Cell 78(1): 137–147.

Özel, M. N., F. Simon, S. Jafari, I. Holguera, Y. C. Chen, N. Benhra, R. N. El-Danaf, K. Kapuralin, J. A. Malin, N. Konstantinides and C. Desplan (2021). “Neuronal diversity and convergence in a visual system developmental atlas.” Nature 589(7840): 88–95.

Pan, D. and G. M. Rubin (1998). “Targeted expression of teashirt induces ectopic eyes in Drosophila.” Proceedings of the National Academy of Sciences 95(26): 15508–15512.

Pappu, K. S., E. J. Ostrin, B. W. Middlebrooks, B. T. Sili, R. Chen, M. R. Atkins, R. Gibbs and G. Mardon (2005). “Dual regulation and redundant function of two eye-specific enhancers of the Drosophila retinal determination gene dachshund.”

Prüßing, K., A. Voigt and J. B. Schulz (2013). “Drosophila melanogaster as a model organism for Alzheimer’s disease.” Molecular Neurodegeneration 8(1): 35.

Quiquand, M., G. Rimesso, N. Qiao, S. Suo, C. Zhao, M. Slattery, K. P. White, J. J. Han and N. E. Baker (2021). “New regulators of Drosophila eye development identified from temporal transcriptome changes.” Genetics 217(4): iyab007.

Samson, M.-L. and F. Chalvet (2003). “found in neurons, a third member of the Drosophila elav gene family, encodes a neuronal protein and interacts with elav.” Mechanisms of development 120(3): 373–383.

Sang, T.-K. and G. R. Jackson (2005). “Drosophila models of neurodegenerative disease.” NeuroRx 2: 438–446.

Seimiya, M. and W. J. Gehring (2000). “The Drosophila homeobox gene optix is capable of inducing ectopic eyes by an eyeless-independent mechanism.” Development 127(9): 1879–1886.

Shapiro, E., T. Biezuner and S. Linnarsson (2013). “Single-cell sequencing-based technologies will revolutionize whole-organism science.” Nature Reviews Genetics 14(9): 618–630.

St Pierre, S. E., M. I. Galindo, J. P. Couso and S. Thor (2002). “Control of Drosophila imaginal disc development by rotund and roughened eye: differentially expressed transcripts of the same gene encoding functionally distinct zinc finger proteins.”

Stultz, B. G., H. Lee, K. Ramon and D. A. Hursh (2006). “Decapentaplegic head capsule mutations disrupt novel peripodial expression controlling the morphogenesis of the Drosophila ventral head.” Developmental biology 296(2): 329–339.

Su, T. T. and P. H. O’Farrell (1997). “Chromosome association of minichromosome maintenance proteins in Drosophila mitotic cycles.” The Journal of cell biology 139(1): 13–21.

Su, T. T. and P. H. O’Farrell (1998). “Chromosome association of minichromosome maintenance proteins in Drosophila endoreplication cycles.” The Journal of cell biology 140(3): 451–460.

Taylor, S. R., G. Santpere, M. Reilly, L. Glenwinkel, A. Poff, R. McWhirter, C. Xu, A. Weinreb, M. Basavaraju and S. J. Cook (2019). “Expression profiling of the mature C. elegans nervous system by single-cell RNA-Sequencing.” BioRxiv: 737577.

Wawersik, S. and R. L. Maas (2000). “Vertebrate eye development as modeled in Drosophila.” Human Molecular Genetics 9(6): 917–925.

Wen, L., G. Li, T. Huang, W. Geng, H. Pei, J. Yang, M. Zhu, P. Zhang, R. Hou and G. Tian (2022). “Single-cell technologies: From research to application.” The Innovation 3(6).

Xiong, W.-C., H. Okano, N. H. Patel, J. A. Blendy and C. Montell (1994). “repo encodes a glial-specific homeo domain protein required in the Drosophila nervous system.” Genes & development 8(8): 981–994.

Yamaguchi, M. and H. Yoshida (2018). Drosophila as a Model Organism. Drosophila Models for Human Diseases. M. Yamaguchi. Singapore, Springer Singapore: 1–10.

Yeung, K., K. K. Bollepogu Raja, Y.-K. Shim, Y. Li, R. Chen and G. Mardon (2022). “Single cell RNA sequencing of the adult Drosophila eye reveals distinct clusters and novel marker genes for all major cell types.” Communications Biology 5(1): 1370.

